# Targeting CBP/p300 Overcomes Acquired Vincristine Resistance in Medulloblastoma

**DOI:** 10.64898/2026.07.28.740967

**Authors:** Goktug Karabiyik, Ozlem Yedier-Bayram, Filiz Senbabaoglu Aksu, Tolga Lokumcu, Ali Cenk Aksu, Fidan Seker-Polat, Ezgi Ozyerli-Goknar, Adam P Cribbs, Udo Oppermann, Tugba Bagci-Onder

## Abstract

**Background:** Medulloblastoma is the most common malignant pediatric brain tumor. Although advances in conventional therapies have improved survival over the years, acquired drug resistance remains a major barrier to durable cure. As dysregulation of epigenetic mechanisms is increasingly recognized as a driver of medulloblastoma pathogenesis and therapeutic adaptation, targeting epigenetic vulnerabilities represents a promising strategy to overcome treatment resistance.

**Methods:** We generated vincristine-resistant medulloblastoma cell line models and performed chemical screening to identify therapeutically targetable vulnerabilities. Candidate hits were validated using transcriptomic analyses, chromatin immunoprecipitation, and CRISPR-mediated genetic ablation to define the molecular mechanisms underlying drug sensitivity.

**Results:** Chemical screening identified multiple active epigenetic compound classes capable of resensitizing vincristine-resistant medulloblastoma cells, including histone methyltransferase inhibitors, histone deacetylase inhibitors, and bromodomain inhibitors. Among these, the CBP/p300 bromodomain inhibitor SGC-CBP30 emerged as the most potent sensitizer to vincristine. Transcriptomic profiling revealed that, while *ABCB1* was among the most highly upregulated genes in resistant cells, SGC-CBP30 treatment selectively downregulated *ABCC3* and *ABCA4*, an effect not observed in parental cells. Mechanistically, chromatin immunoprecipitation demonstrated enrichment of p300 and H3K27ac at the *ABCC3* and *ABCA4* promoters in resistant cells, which was markedly reduced following SGC-CBP30 treatment. Consistent with these findings, genetic ablation of CREBBP or EP300 phenocopied the effects of pharmacological inhibition. Analysis of patient datasets further demonstrated elevated *CREBBP*, *EP300*, and *ABCC3* expression in SHH MB, with positive correlations between *ABCC3* and both *CREBBP* and *EP300*, supporting the clinical relevance of this regulatory axis.

**Conclusions:** Together, our findings demonstrate that CBP/p300 activity contributes to acquired vincristine-resistance in medulloblastoma. Targeting this axis represents a promising strategy to overcome drug resistance and enhance the efficacy of vincristine-based chemotherapy particularly in the context of relapsed or refractory disease.

**PLAIN ENGLISH SUMMARY:** Medulloblastoma is the most common cancerous brain tumor in children. Although many children respond well to the treatment, some tumors become resistant to chemotherapy, making them much harder to treat. Understanding why this resistance develops could lead to better treatment options for children whose cancer returns or no longer respond to therapy.

In this study, we created laboratory models of medulloblastoma that had become resistant to the chemotherapy drug vincristine. We then tested a collection of drugs to identify compounds, which would restore the cancer cells’ sensitivity to treatment. We have discovered that several drugs were effective, with one compound, called SGC-CBP30, showing particularly strong activity.

We investigated how SGC-CBP30 works and found that it decreases the activity of genes that are linked to chemotherapy resistance. Using multiple complementary experimental approaches, we confirmed that this gene-regulating pathway plays an important role in helping medulloblastoma cells survive treatment.

Our findings suggest that targeting this pathway could restore the effectiveness of chemotherapy in drug-resistant tumors. Although further research is needed before this approach can be used in patients, these results provide a promising foundation for developing new treatments for children with relapsed or treatment-resistant medulloblastoma.

## BACKGROUND

Cancer remains a major global health burden and is the second leading cause of mortality worldwide^1^. Medulloblastoma (MB), the most common malignant brain tumor in children, accounting around 20% of all cases, arises in the cerebellum or posterior fossa and is classified by the WHO as a grade IV embryonal tumor due to its high proliferative and metastatic potential^2^. Current treatment for MB largely relies on maximal surgical resection followed by craniospinal irradiation and chemotherapy with recent advances focusing on improved risk stratification and the development of molecularly targeted therapies^3–5^. Despite these advances, chemotherapy remains a cornerstone of MB treatment. Chemotherapeutic agents such as vincristine, cisplatin, carboplatin, and lomustine are widely used in the treatment of a broad range of pediatric and adult malignancies. Among these, vincristine, a vinca alkaloid that disrupts microtubule polymerization and induces mitotic arrest, is a key component of treatment regimens for MB, as well as neuroblastoma, leukemias, lymphomas, and nephroblastoma^6^. However, the clinical benefit of vincristine is often limited by dose-limiting toxicities, particularly peripheral neuropathy, and by the development of acquired drug resistance, especially in relapsed disease^7^.

Although chemotherapeutic agents have remarkably improved the overall survival of patients with MB over the past decades, the emergence of drug resistance remains a major barrier to effective long-term treatment. Cancer cells can acquire resistance to chemotherapeutic agents through multiple mechanisms, including enhanced drug efflux, reduced drug uptake, alterations in drug targets and activation of pro-survival signaling pathways^8^. Among these, active drug efflux is one of the best-characterized mechanisms, whereby membrane transporters reduce intracellular concentrations of chemotherapeutic agents and thereby diminish their efficacy^9^. While there are several transporters that take part in these procedures, responsible for different chemicals, the ATP-binding Cassette (ABC) transporter superfamily plays a particularly important role^10^. ABCB1 (P-glycoprotein) is especially important in MB, as it is responsible for the efflux of several chemotherapeutic agents, including vincristine^11^. Consequently, elevated expression of ABCB1 can drive multidrug resistance (MDR), whereby resistance to a single ABCB1 substrate confers cross-resistance to multiple drugs, ultimately limiting the effectiveness of standard chemotherapy^8^.

Growing evidence indicates that epigenetic alterations promote tumor heterogeneity and cellular plasticity, thereby contributing to key malignant phenotypes, including drug resistance. Given the complex and multifactorial nature of therapy resistance, targeting a single downstream effector, such as ABCB1, is unlikely to be sufficient to completely overcome resistance^12^. Instead, targeting upstream epigenetic regulators that orchestrate broad transcriptional programs may provide a more effective therapeutic strategy. Previous studies showed that dysregulation of chromatin-associated pathways contributes to both tumor initiation and the acquisition of resistance across solid and hematological malignancies^13^. In MB, genomic studies have further revealed that, in addition to established driver genes, approximately 50% of tumors harbour at least one somatic mutation in a chromatin-modifying gene^14–17^. In line with these, several drugs targeting epigenetic modifiers are undergoing clinical evaluation, with some already entering clinical practice^18–21^.

Bromodomains (BRDs) are evolutionarily conserved protein interaction modules that recognize acetylated lysine residues on histones and other proteins. Although they lack intrinsic enzymatic activity, bromodomain-containing proteins regulate gene expression by facilitating chromatin remodelling, recruiting transcriptional regulatory complexes, and functioning as transcriptional co-activators or co-repressors^22^. Among these, the closely related histone acetyltransferases CBP and p300 (also known as KAT3A and KAT3B, respectively), each contain a single bromodomain motif and share 96% sequence similarity^23^.

As master transcriptional co-activators, CBP and p300, regulate diverse biological processes including genome maintenance, development, cell proliferation and oncogenic transformation, making them attractive therapeutic targets^24–26^.

In this study, we established models of acquired vincristine resistance in MB and performed an epigenetic compound screen to identify regulators of drug resistance. We demonstrated that pharmacological inhibition of the CBP/p300 bromodomain resensitizes vincristine-resistant cells by suppressing the expression of ABC transporters ABCC3 and ABCA4. Finally, analysis of patient datasets supports the clinical relevance of this regulatory axis, identifying CBP/p300 as a therapeutically actionable epigenetic dependency in acquired vincristine resistance.

## METHODS

### Cell culture and generation of vincristine-resistant cell lines

Human medulloblastoma cell line Daoy (HTB-186) and human embryonic kidney 293T (CRL- 3216) cells, were purchased from American Tissue Type Culture Collection (ATCC); human astrocytes were purchased from ScienCell, USA. Daoy, HEK-293T, and astrocyte cells were cultured in Dulbecco’s Modified Eagle Medium (DMEM; Gibco, USA) supplemented with 10% fetal bovine serum (FBS; Gibco, USA) and 1% Penicillin-Streptomycin (Gibco, USA). All cells were grown and maintained at 37 °C in a humidified incubator with 5% CO2. All cell lines were routinely screened for mycoplasma contamination using MycoAlert Mycoplasma Detection Kit (Lonza, CH).

Vincristine-resistant cells were established using a dose escalation protocol. Parental Daoy cells were initially treated with two different concentrations of vincristine (1 nM and 2 nM) for a period of three months. Vincristine concentrations were then gradually increased to a final concentration of 10 nM and 8 nM respectively. Age matching cells were maintained in a drug-free medium throughout the experiment. After eight months of continuous vincristine exposure, two vincristine resistant cell lines were generated namely, V1-10 and V2-8, based on their respective initial and final vincristine concentrations. Cells were maintained at 10 nM or 8 nM, respectively during subculturing, unless mentioned “drug holiday".

### Reagents

Vincristine (#11764) was purchased from Cayman Chemicals, USA. An epigenetic probe library consisting of 89 inhibitors targeting chromatin-modifying proteins **(Supplementary Table 1)** was provided by Prof. Udo Oppermann (Structural Genomics Consortium, University of Oxford, UK). All individual drugs, including SGC-CBP30, A485, CBP/BRD4, I-CBP112, PFI- 4, OF-1, NVS-CECR2 were supplied from Selleckchem, USA.

### Cell viability assays

Cells were seeded at a density of 2,000 cells/well into black 96-well plates (Corning, USA) and treated with the indicated probes and/or vincristine the following day. After 72 hours of treatment, cell viability was assessed using CellTiter-Glo Luminescent Cell Viability Assay (Promega, USA) according to the manufacturer’s instructions. Each treatment condition was tested in triplicates. Viability was calculated by normalizing luminescence signals to those of untreated control wells.

### Live cell microscopy

Live-cell imaging was performed using a Leica DMI8 inverted microscope (Zeiss, Germany) with 10X air objective in a chamber at 37°C with 5% CO2. Time-lapse images were acquired every 5 minutes over a 24-hour (Figure 1) or 32-hour (Figure 3) period. Quantification of normalized cell numbers per frame was conducted using ImageJ software^27^, based on at least three randomly selected fields per condition at each time point. All videos are supplied as **Additional Files 1-12.**

**Figure 1.**
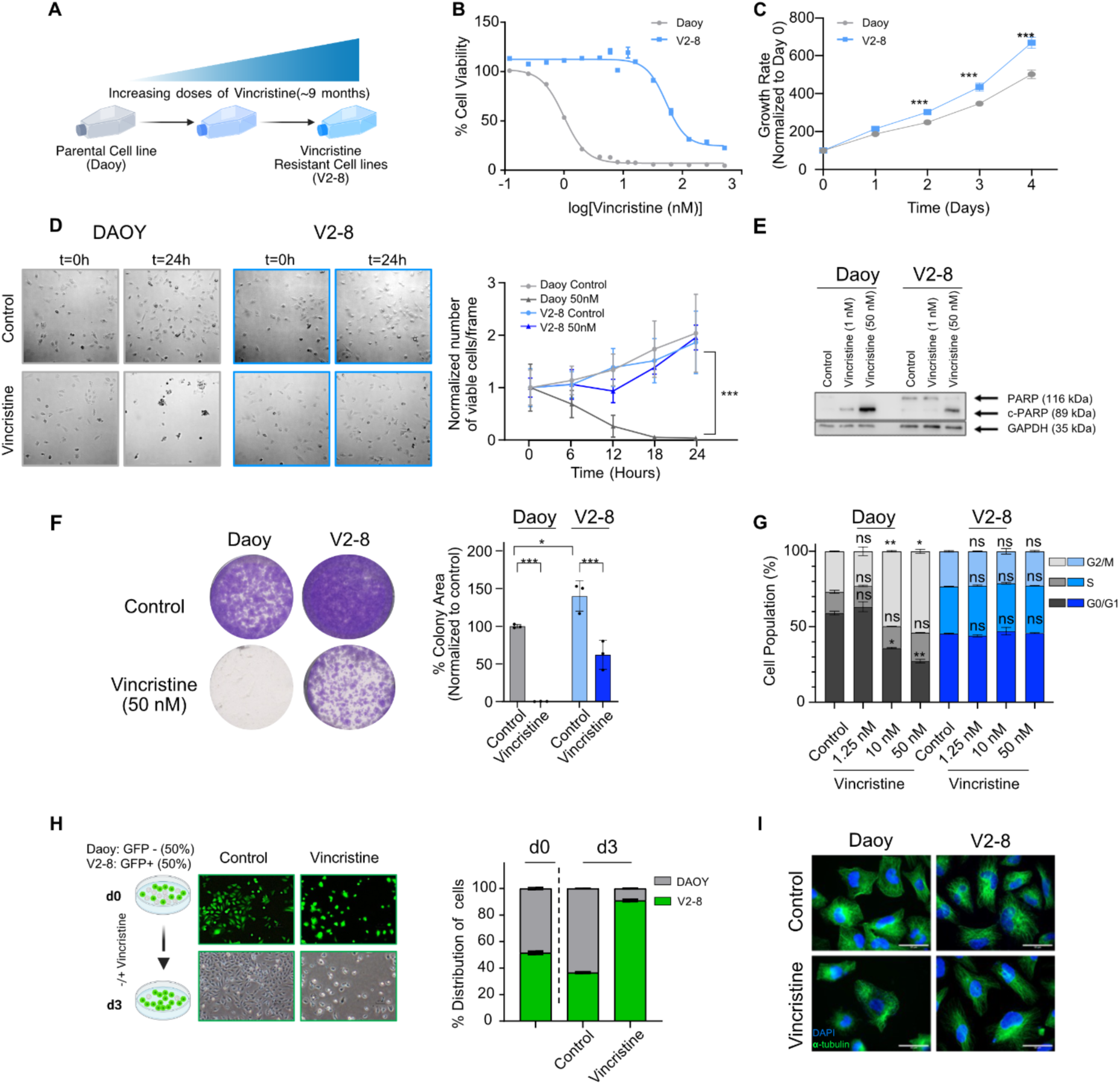
Generation and characterization of vincristine-resistant medulloblastoma cell lines. A) Schematic representation of the establishment of vincristine-resistant cell lines using a dose-escalation approach. B) Dose-response curves of Daoy and V2-8 cell lines treated with vincristine. C) Proliferation curves Daoy and V2-8 cell lines. D) Live-cell imaging of Vincristine-treated cells over 24 hours. Representative images and quantification of viable cells per frame are shown. Compared to Daoy-50 nM, all groups were shown to be significantly different (***p<0.001) while no statistical significance was observed in between other comparisons. E) Western blot analysis of PARP cleavage in Daoy and V2-8 cells treated with 1 nM or 50 nM vincristine for 24 hours. F) Long-term clonogenic assay on parental and resistant cell lines treated with 50 nM vincristine and quantification of the area covered by colonies. G) Cell cycle assay after 16 hours of DMSO or vincristine treatment (1.25, 10, 50 nM). H) Growth competition assay between Daoy and V2-8-GFP cells in the presence of DMSO or vincristine and its quantification. I) Immunofluorescence staining of alpha-Tubulin and DAPI in Daoy and V2-8 cells treated with vincristine for 24 hours. *p < 0.05, **p < 0.01, ***p < 0.001. Unless otherwise specified, vincristine was used at 1.25 nM for Daoy cells and 50 nM for V2-8 cells.

### Immunofluorescence staining

Cells were seeded on coverslips in 24-well plates at a density of 20,000 cells per well and treated the next day with the indicated compounds alone or in combination with vincristine. After 24 hours of incubation, cells were washed with PBS, fixed with 4% PFA for 5 minutes, permeabilized with 0.1% Triton X-100 and blocked for 15 minutes. Coverslips were incubated with primary antibodies **(Supplementary Table 2)** overnight at 4°C followed by secondary antibody incubation for 1 hour at room temperature in the dark. Images were acquired at 40X magnification using a Zeiss Axio Imager M1 (Zeiss, Germany).

### Epigenetic probes library screen

Cells were seeded at a density of 2,000 cells per well in black 96-well plates. The next day, cells were treated in triplicates with epigenetic probes at 1:1000 dilution from the epigenetic probe library alone **(Supplementary Table 1)** or in combination with vincristine at a concentration of 1.2 nM for Daoy and 50 nM for V2-8, corresponding to the IC50 values previously determined for each respective cell line under the same assay conditions. After 72 hours of treatment, cell viability was measured using CellTiter-Glo Luminescent Cell Viability Assay as described above. A threshold of 50% cell viability relative to untreated controls was used to define hits. Hits were excluded if their inactive probe controls also affected cell viability.

### Western blot

Whole cell extracts were prepared using lysis buffer containing 1% NP-40, 150 mM NaCl, 1mM EDTA, 50 mM Tris-HCl (pH 7.8), 1 mM NaF, 0.5 mM PMSF, and 1X protease inhibitor cocktail (Complete Protease Inhibitor Cocktail Tablets, Roche). The protein concentrations were determined using the Pierce BCA Protein Assay Kit (Thermo Fisher Scientific, USA) according to the manufacturer’s instructions. 50 µg protein per sample was run on SDS-PAGE and transferred onto Immun-Blot PVDF Membranes (Bio-Rad, USA) using the Trans-Blot Turbo Transfer System (Bio-Rad, USA). Antibodies and their respective dilutions are listed in **Supplementary Table 2**.

### Annexin V/Dead cell assay

Annexin V/Dead cell staining was performed using Muse Annexin V & Dead Cell kit (Luminex, USA) according to manufacturer’s instructions. Briefly, cells were collected, washed with cold PBS containing 1% FBS, and resuspended in the kit’s staining buffer. Cells were then incubated with Annexin V & Dead Cell reagent for 20 minutes at room temperature and analyzed using Muse Cell Analyzer (Merck, USA) with 5,000 events per sample. Gates were set based on parental cell untreated controls.

### YO-PRO-1 staining

Cells were seeded to 6-well plates at a density of 50,000 cells per well. The following day, cells were treated with vincristine and 1 μM SGC-CBP30. After 72 hours of incubation, cells were stained with 1 μM YO-PRO-1 (Invitrogen:Y3603, Thermo Fisher, USA) and 1:1000 PI (1 mg/ml) for 15 minutes at 37 °C in the dark. Fluorescence images were acquired using a Nikon Eclipse TS100 Inverted Fluorescence Microscope. Quantification of YO-PRO-1 positive cells was performed using ImageJ software^27^.

### Long-term clonogenic assay

Long-term clonogenic assay was performed to assess the long-term impact of drug exposure on cell fitness and colony formation. Briefly, 5,000 cells per well were seeded in triplicate in 6- well plates (Corning, USA). The following day, cells were treated with the indicated vincristine and/or SGC-CBP30 and A485 doses. Following three days of drug treatment, media were replaced and cells were incubated until colonies formed in control wells. Plates were then washed with PBS, fixed with ice-cold 100% methanol for 5 minutes, and stained with crystal violet for 15 minutes. After removing excess stain, plates were dried overnight and imaged against a white background. Colony area was quantified using the ImageJ^27^ and normalized to controls.

### RNA sequencing

Total RNA was isolated from cells treated with 1 μM SGC-CBP30 for 72 hours and untreated controls using the NucleoSpin RNA kit (Macherey-Nagel, Germany). RNA quality was assessed on a TapeStation (Agilent, USA), with RIN values ranging from 7.5 to 9.5. Libraries were prepared using the NEBNext Ultra RNA kit with TruSeq indexes and sequenced on an Illumina NextSeq 500 (paired-end, 2 × 80 bp) to an average depth of 112 million reads per sample (range: 47–168 million).

Reads were aligned to the human genome (hg38) using STAR, and transcript quantification was performed with Salmon. Quality control was conducted using FastQC. Gene-level counts were generated with tximport, and differential expression analysis was performed using DESeq2. Differentially expressed genes (DEGs) were defined as |log₂ fold change| > 1 and adjusted p-value < 0.05. Gene set enrichment analysis (GSEA) was performed using fgsea on a pre-ranked gene list based on log₂ fold change, testing MSigDB Hallmark, KEGG, and Reactome gene sets (FDR < 0.05)^28^.

### Quantitative RT-PCR

Total RNA was isolated using the NucleoSpin RNA isolation kit (Macherey-Nagel) according to the manufacturer’s instructions. For each sample 1000 ng of total RNA was used to synthesize cDNA using M-MLV Reverse Transcriptase (Invitrogen) with manufacturer’s instructions. qRT-PCR was performed using LightCycler® 480 SYBR Green I Master mix (Roche, USA) on a LightCycler 480 Instrument (Roche, USA). Each condition was tested in triplicates. Relative gene expression levels were calculated using the 2^(-ΔΔCt)^ method. Primer sequences are provided in **Supplementary Table 3**.

### Calcein-AM assay

Cells were seeded in 24-well plates at a density of 5,000 cells per well. The following day, cells were treated with indicated drugs for 24 hours. Cells were then incubated with 125 nM Calcein-AM (Thermo Fisher, USA), for 30 minutes at 37°C and green fluorescence signal was imaged using Cytation 5 (Biotek, USA) and quantified with a Synergy H1 Plate reader (Biotek, USA).

### CHIP-qPCR

ChIP-qPCR was performed to examine the H3K27ac and p300 occupancy at the promoter of selected ABC transporter genes. 2 x 10^6^ cells were crosslinked with 1 % formaldehyde at room temperature for 10 minutes. Then, quenched with 0.125 M glycine. Cells were washed with ice-cold PBS and lysed with ChIP lysis buffer, followed by sonication with Bioruptor (Diagenode, Belgium). Sonicated samples were centrifuged at top speed for 10 minutes to remove the debris, then supernatant was 1:10 diluted in ChIP dilution buffer, and 10% of the diluted sample was used as input.

Samples were incubated overnight with 5 µg of target antibody (Anti-H3K27ac, Anti-p300, or non-specific IgG, **Supplementary Table 2**) at +4°C followed by protein A/G bead binding for two hours at room temperature. Then, beads were sequentially washed with low-salt, high-salt, LiCl and TE buffers. Finally, chromatin was eluted and reverse crosslinking was applied with 0.2 M NaCl overnight at 65°C. DNA was purified with a Qiagen PCR purification kit (Qiagen, Germany), and enrichment at target promoter regions was analyzed by qRT-PCR **(Supplementary Table 4)**.

### CRISPR/Cas9 knockout

Guide RNAs (sgRNAs) targeting *CREBB (*CBP), *EP300 (*p300) and Non-Targeting1 (NT1) were cloned into pLentiGuideV2 plasmid, as described^29^. Oligonucleotide sequences used for cloning are listed in **Supplementary Table 5**. Briefly, lentiviruses for sgRNA containing LentiGuide plasmids and LentiCas9-Blast (Addgene, #52962) were produced using FuGENE 6 transfection reagent (Promega, USA) according to manufacturer’s recommendations. Cells were transduced with Cas9 lentiviruses at MOI∼10 and selected with 10 µg/ml blasticidin for 6-7 days. Cas9-stable Daoy, V1-10 and V2-8 cells were subsequently transduced with sgRNA-containing lentiviruses at MOI∼10 and selected with puromycin (2 µg/ml for Daoy; 10 µg/ml for V1-10 and V2-8) for 3-4 days. At day 10 post-sgRNA transduction, cells were seeded for clonogenic assay and processed as described above.

### Patient data analysis

Publicly available data were retrieved from the Gliovis platform (https://gliovis.bioinfo.cnio.es). Gene expression datasets from Northcott et al.^30^ (GSE37385) were preprocessed and normalized by the Gliovis platform using its standard processing pipeline, and the processed expression values were exported and analyzed to generate mRNA expression comparisons between molecular groups, gene–gene correlation analyses, and a heatmap of selected genes.

### Statistical analysis

Statistical analysis was conducted using Graphpad Prism (v 8.0), USA. All normalizations were performed to untreated control cells. Comparisons between two groups were evaluated using a two-tailed Student’s t-test. For experiments comparing multiple groups across a single variable, a one-way ANOVA while for experiments involving two independent variables a two-way ANOVA followed by Tukey’s post hoc multiple comparisons test was utilized. Significance thresholds were defined as *p<0.05, **p<0.01, and ***p<0.001.

## RESULTS

### Establishment of vincristine-resistant cells

To model acquired drug resistance in MB, Daoy cells were exposed to escalating concentrations of vincristine over eight months, generating two vincristine-resistant cell lines V2-8 and V1-10, named according to initial and final vincristine concentrations (nM) used during selection (**Figure 1.A, S1.A**). Cells were subsequently maintained at their respective final vincristine doses throughout the study. Compared with age-matched parental (naïve) cells, V2-8 and V1-10 cells exhibited approximately 50-fold and 100-fold increases in vincristine resistance, respectively (**Figure 1.B, Figure S1.B**). Importantly, following a 50-day drug “holiday”, V2-8 cells retained a stable IC50, confirming maintenance of the acquired resistance phenotype (**Figure S2.A**). Both resistant lines displayed a significant increase in proliferation rates compared with parental cells (**Figure 1.C, S1.C**). Live cell imaging demonstrated that, whereas parental Daoy cells rapidly ceased proliferating and progressively disappeared following vincristine treatment, V2-8 cells continued to proliferate despite continuous drug exposure for 32 hours (**Figure 1.D**). Supporting this, vincristine-induced PARP cleavage was markedly attenuated in resistant cells (**Figure 1.E**). Consistently, long-term colony formation assay showed that V2-8 and V1-10 cells survived 50 nM of vincristine whereas the parental cells were depleted (**Figure 1.F, S1.D**). Moreover, unlike parental cells that accumulated in G2/M following vincristine exposure, resistant cells largely escaped vincristine-induced mitotic arrest (**Figure 1.G, S1.E**). To further validate the resistant phenotype, GFP-expressing V2-8 cells were co-cultured with age-matched parental Daoy cells. After 72 hours under control conditions, the proportion of GFP-positive V2-8 cells showed a slight decrease, likely reflecting a slower proliferation rate in co-culture. In contrast, vincristine treatment selectively depleted the GFP-negative parental population, resulting in a marked enrichment of GFP-positive V2-8 cells and further confirming their resistance to vincristine (**Figure 1.H**). Similarly, α-tubulin immunofluorescence after 24 hours of vincristine treatment showed microtubule disruption in the parental cells while V2-8 cells largely preserved microtubule organization (**Figure 1.I**).

### Epigenetic probe library screening identifies potential therapeutic candidates

To identify epigenetic vulnerabilities associated with acquired vincristine resistance, we performed chemical screens using an epigenetic probe library in parental Daoy and vincristine-resistant V2-8 cells, both in the absence and presence of vincristine (**Figure 2.A-B**). Several probes, including histone deacetylase inhibitors (eg. belinostat, romidepsin) and histone demethylase inhibitors (eg. KDOBA67, IOX-1) reduced viability in both cell lines, whereas Trichostatin A and GSK-J4 displayed selective toxicity toward the V2-8 cells, indicating distinct epigenetic dependencies (**Figure 2.C-D**). More importantly, multiple epigenetic probes selectively sensitized V2-8 cells to vincristine, with bromodomain inhibitors emerging as the most prominent class of hits. In particular, inhibitors targeting the CBP/p300 and BRD protein families (CBP/BRD4, SGC-CBP30, I-CBP112, PFI-4, OF-1, and NVS-CECR2) markedly potentiated vincristine-induced cytotoxicity in V2-8 cells while exerting minimal effects on parental cells, either alone or in combination with vincristine (**Figure 2.E-G**). To validate these findings, representative bromodomain inhibitors were evaluated in independent combination experiments. All tested compounds significantly enhanced vincristine sensitivity in V2-8 cells with minimal effects on parental cells **(Figure 2.H).** Moreover, although V1-10 cells were not included in the primary screen, bromodomain inhibitors similarly restored vincristine sensitivity in this independently generated resistant model **(Figure S1.F).**

**Figure 2.**
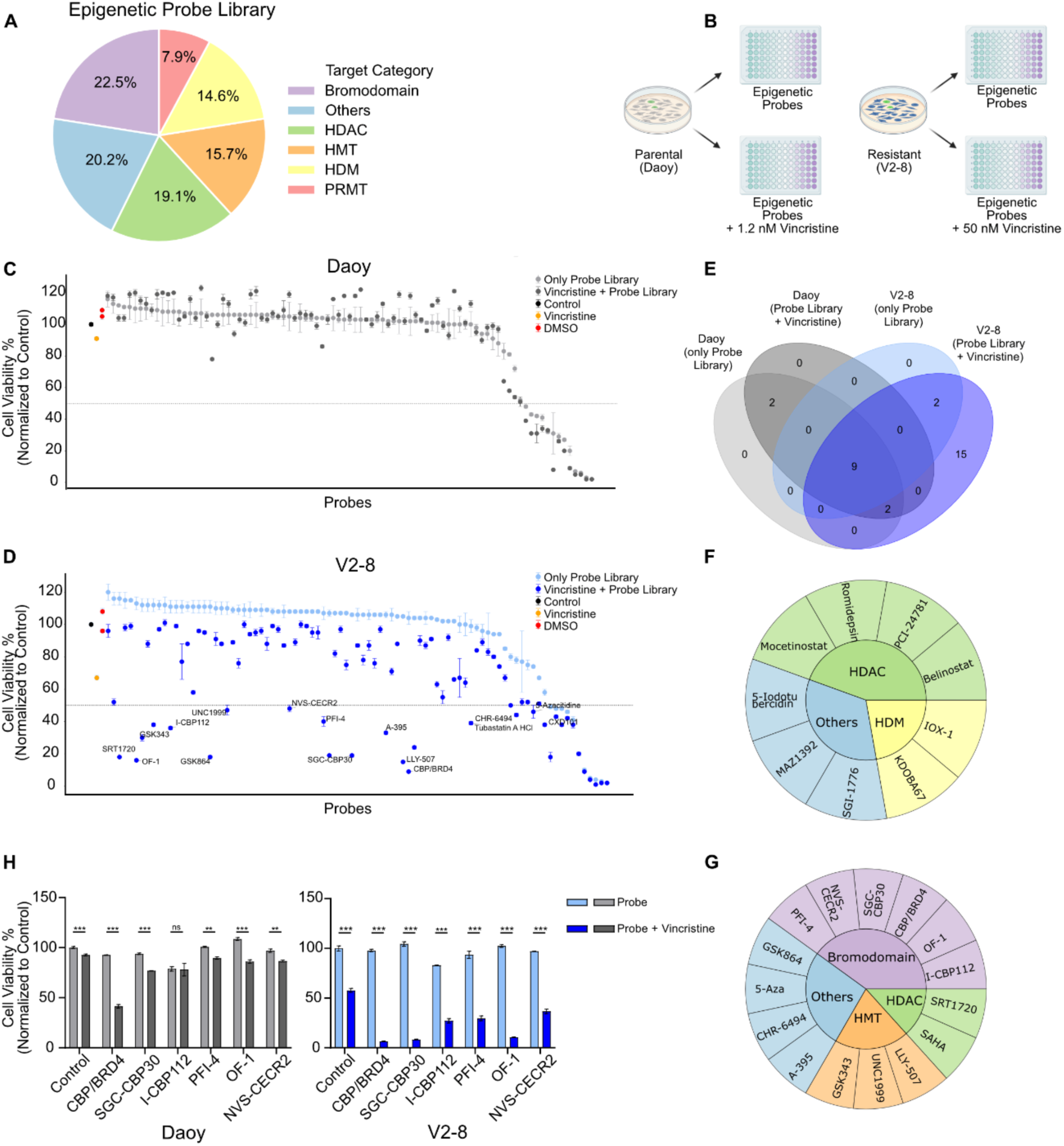
Epigenetic probe library screening identifies potential therapeutic candidates. A) Overview of targets included in the epigenetic probe library. HDAC: Histone Deacetylase, HMT: Histone Methyltransferase, HDM: Histone Demethylase, PRMT: Protein Arginine Methyltransferase B) Schematic representation of the screening workflow. C,D) Results of the epigenetic probe screen conducted on Daoy and V2-8 cell lines. A threshold of 50% cell viability relative to untreated controls was used to define hits. E) Venn diagram summarizing screen hits. F) Hits showing toxicity towards both cell lines. G) Hits that sensitize only the V2-8 cell line to vincristine. H) Validation of selected bromodomain inhibitors in parental Daoy and V2-8 cell lines. Statistical analysis was performed using one way ANOVA. *p < 0.05, **p < 0.01, ***p < 0.001, ns: not significant.

### Bromodomain inhibition restores vincristine sensitivity in resistant MB cells

SGC-CBP30 is a selective bromodomain inhibitor targeting CBP and p300^31^. We further evaluated its potential sensitization effect on both V2-8 and parental cells (**Figure 3.A**). SGC- CBP30 sensitized V2-8 cells to vincristine in a dose-dependent manner, with 1 μM significantly enhancing vincristine sensitivity and 5 μM restoring the IC50 to levels comparable to those of parental cells (**Figure 3.B**). We observed a similar sensitization effect in the independently generated V1-10 cells when vincristine was combined with SGC-CBP30 (**Figure S1.G**).

**Figure 3.**
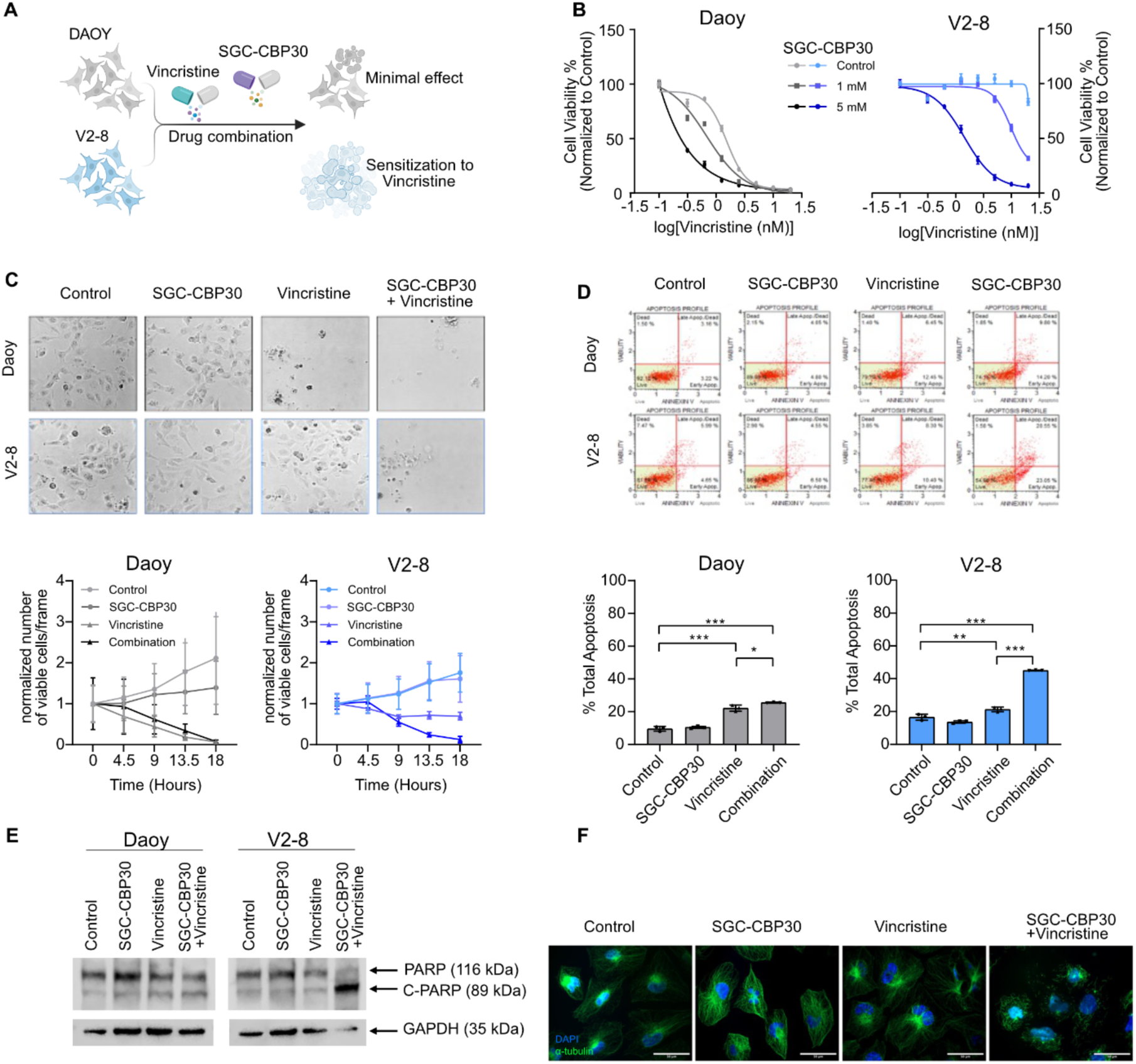
Bromodomain inhibition restores vincristine sensitivity in resistant MB cells. A) Schematic representation of combination treatment. B) Dose-response curves for vincristine and SGC-CBP30 treatment in Daoy and V2-8 cells. C) Representative snapshots of live-cell imaging of vincristine and SGC-CBP30 treatment over 32 hours (*top*). Three frames per condition were quantified up to 18 hours (*bottom*). D) Annexin V/Dead cell assay following treatment with SGC-CBP30 and/or vincristine for 24 hours on Daoy and V2-8 cells. E) Western Blot analysis of PARP, cleaved-PARP (c-PARP) and GAPDH (loading control) on Daoy and V2-8 cells treated with SGC-CBP30 and/or vincristine for 24 hours. F) Immunofluorescence staining of alpha-Tubulin and DAPI in V2-8 cells treated with SGC-CBP30 and/or vincristine for 24 hours. Statistical analysis was performed using two-way ANOVA. *p < 0.05, **p < 0.01, ***p < 0.001, ns: not significant.

Live cell imaging showed that vincristine alone depleted parental cells but had only modest effects on V2-8 cells. In contrast, the combination treatment with SGC-CBP30 and vincristine resulted in rapid depletion of V2-8 cells, further indicating restoration of sensitivity (**Figure 3.C**). Consistent with this finding, Annexin V/Dead cell analysis performed using cell line specific vincristine doses revealed a significant increase in the percentage of apoptotic cells following combination treatment in V2-8 cells (**Figure 3.D**). Likewise, SGC-CBP30 enhanced vincristine-induced PARP cleavage in both V2-8 and V1-10 cells, whereas no comparable increase was observed in parental cells (**Figure 3.E, S1.H**). Finally, α-tubulin immunofluorescence demonstrated that, although vincristine alone failed to disrupt the microtubule network in resistant cells, co-treatment with SGC-CBP30 restored induced microtubule disorganization, consistent with re-sensitization to vincristine (**Figure 3.F**).

### CBP/p300 inhibition suppresses ABC transporter expression and activity in vincristine-resistant cells

To investigate the transcriptional mechanisms underlying CBP/p300-mediated vincristine sensitization, we performed bulk RNA sequencing of parental Daoy and vincristine-resistant V2-8 cells treated with either vehicle or SGC-CBP30. Principal component analysis (PCA) revealed clear segregation of all cell line/treatment groups (**Figure 4.A**). Compared with parental cells, V2-8 cells exhibited extensive transcriptional reprogramming, approximately 1000 genes differentially expressed genes (|LFC|>1 and p<0.05), including marked upregulation of *ABCB1*, a key transporter of vincristine, together with multiple members of the ABC and SLC transporter families (**Figure 4.B**). As expected from inhibition of bromodomains associated with histone acetyltransferases, SGC-CBP30 induced a predominantly repressive transcriptional response, particularly in V2-8 cells (**Figure 4.B-C**). Among the differentially expressed genes, 69 were selectively upregulated in V2-8 and downregulated upon SGC- CBP30 treatment, identifying a candidate transcriptional program associated with acquired vincristine resistance (**Figure 4.D-E**). Notably, members of the ABC transporter family, including ABCC3 and ABCA4, were among the most prominently downregulated genes, and KEGG pathway analysis demonstrated partial normalization of the ABC transporter signature following CBP/p300 inhibition (**Figure 4.F**). Gene ontology (GO) enrichment analysis further revealed significant enrichment of transporter related Biological Process (BP) and Molecular Function (MF) gene sets (**Figure 4.G**).

**Figure 4.**
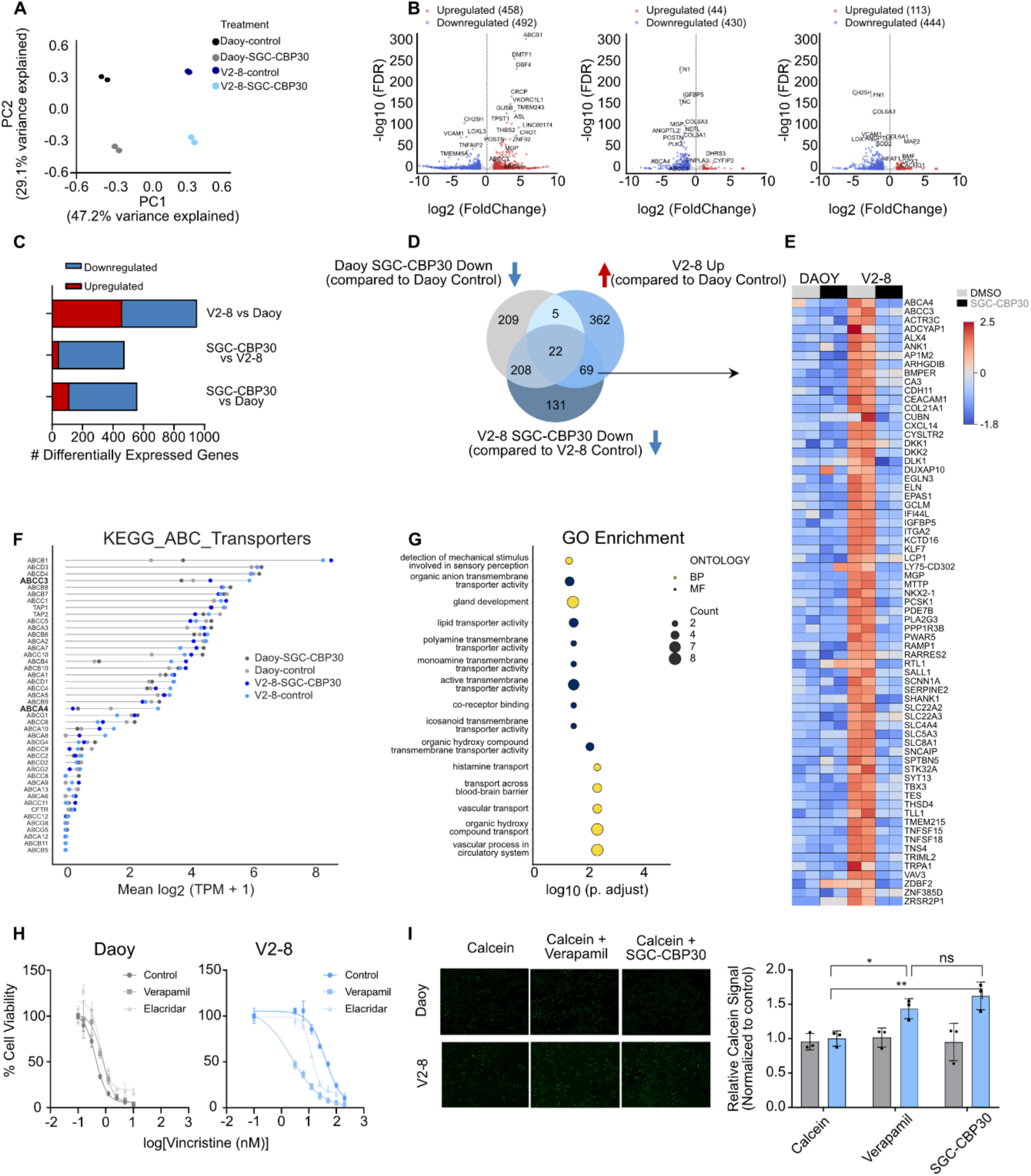
Transcriptomic analysis of Daoy and V2-8 cells treated with SGC-CBP30. A) Principal Component Analysis (PCA) of all samples, showing distinct clustering by treatment and cell line. B) Volcano plots showing differentially expressed genes between, V2-8 vs Daoy, SGC-CBP30-treated vs untreated Daoy, and SGC-CBP30-treated vs untreated V2-8 (LFC>1 and p<0.05). C) Bar graph showing the number of DEGs following SGC-CBP30 treatment. D) Venn diagram illustrating overlap between DEGs in Daoy and V2-8 cells. Genes upregulated in V2-8 and downregulated upon SGC-CBP30 treatment were selected for downstream analysis, including Gene Ontology (GO) enrichment. E) Heatmap of 69 intersecting genes identified above. F) Log₂-transformed TPM values for all treatment groups across genes in the KEGG ABC TRANSPORTERS gene list. G) GO enrichment analysis of the 69 genes identified above. H) Viability assay following treatment with verapamil and elacridar and/or vincristine for 72 hours. I) Calcein-AM uptake assay showing increased intracellular calcein retention in Daoy and V2-8 cells treated with SGC-CBP30 or Verapamil for 72 hours.

To determine whether these transcriptional changes translated into functional inhibition of drug efflux, we first evaluated established MDR inhibitors. Both verapamil and elacridar restored vincristine sensitivity in V2-8 cells, with similar effects observed in the independently generated V1-10 model (**Figure 4.H, S1.I**). Similarly, Calcein-AM accumulation assays demonstrated that SGC-CBP30 increased intracellular calcein retention in V2-8 cells to levels comparable with verapamil treatment, indicating reduced ABC transporter activity following CBP/p300 inhibition (**Figure 4.I**).

### Both bromodomain and acetyltransferase activities of CBP/p300 contribute to acquired vincristine resistance

To further define the molecular mechanisms underlying acquired-vincristine resistance in MB, we compared inhibition of its bromodomain using SGC-CBP30 with inhibition of its histone acetyltransferase (HAT) domain using A485. Unlike SGC-CBP30, which selectively restored vincristine sensitivity in resistant cells, A485 enhanced vincristine-induced cytotoxicity in both parental Daoy and V2-8 cells at their respective vincristine concentrations (**Figure 5.A**). In long-term clonogenic assays, both SGC-CBP30 and A485 markedly potentiated the effects of vincristine in V2-8 cells, although bromodomain inhibition produced a more pronounced reduction in clonogenic survival (**Figure 5.B**). To determine whether the reduced viability reflected increased apoptosis, YO-PRO-1 staining was performed following treatment with subtoxic concentrations of vincristine in combination with SGC-CBP30 or A485. While vincristine alone induced minimal apoptosis in either cell line, both combinations significantly increased apoptotic cell death, with the strongest effect observed following bromodomain inhibition. (**Figure 5.C**). Finally, to genetically validate the role of CBP/p300, *CREBBP* and *EP300* were individually depleted using CRISPR/Cas9. Knockout of either gene significantly sensitized V2-8 cells to vincristine, although the magnitude of sensitization was less pronounced than that achieved by pharmacological inhibition of either the CBP/p300 bromodomain or HAT domain (**Figure 5.D-E**).

**Figure 5.**
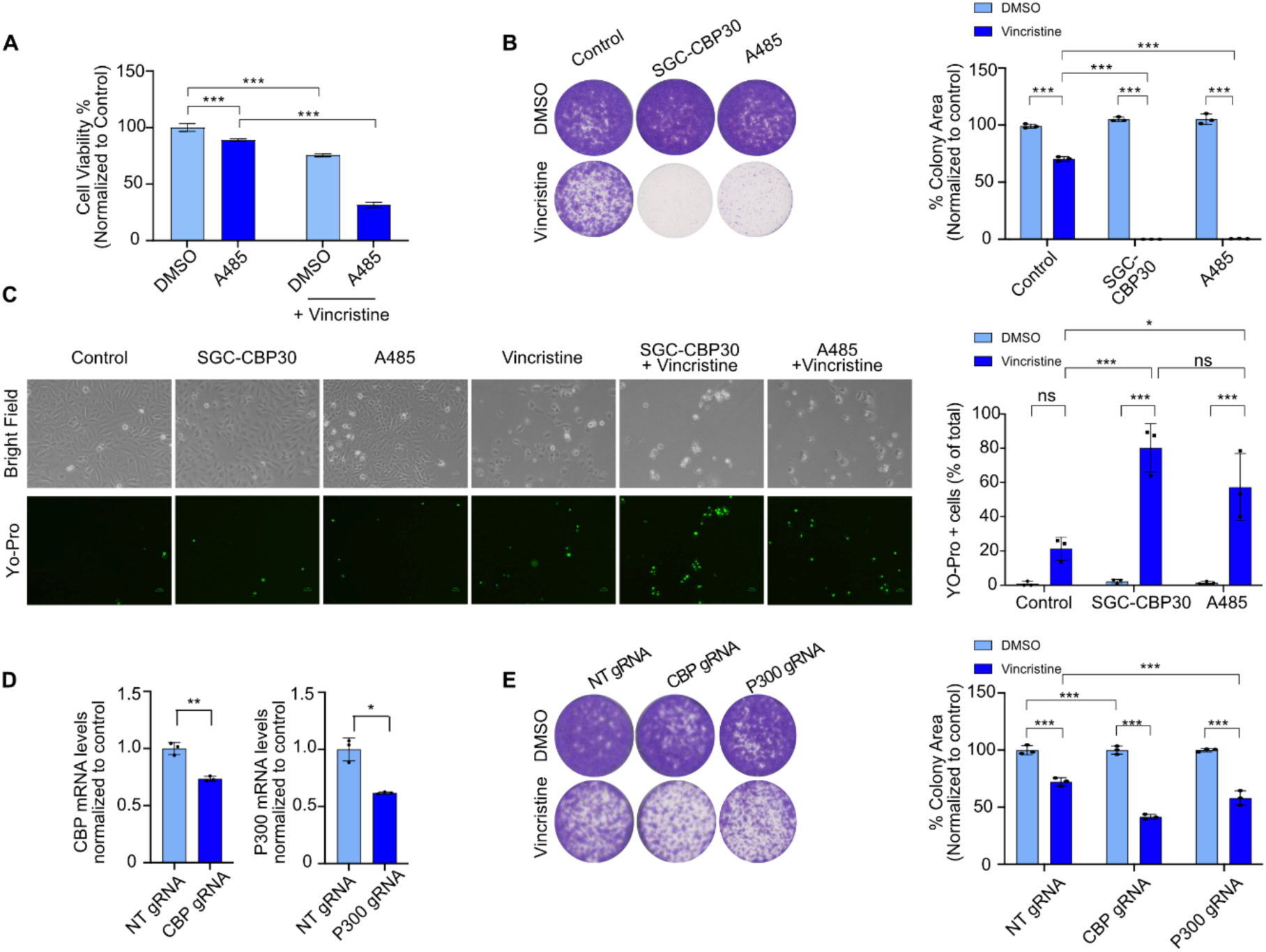
Acquired vincristine resistance is dependent on CBP/P300. A) Cell viability assay on V2-8 cells treated with A485 and/or vincristine for 72 hours. B) Long term clonogenic assay following treatment with SGC-CBP30, A485 and vincristine on Daoy and V2-8 cell lines and quantification of the area covered by colonies. C) YO-PRO-1 staining of V2-8 cells treated with SGC-CBP30 in combination with Vincristine to assess apoptotic cell death. D) *CBP* and *p300* mRNA expression levels upon knockout of respective gens in V2-8 cells. E) Long term clonogenic assay showing the effects of CBP and P300 knockouts in combination with vincristine treatment on V2-8 cells and quantification of the area covered by colonies.

### CBP/p300 directly regulates ABC transporter expression in vincristine-resistant cells

To elucidate whether CBP/p300 directly regulates the expression of ABC transporters associated with acquired vincristine resistance, we performed Chromatin immunoprecipitation (Chip)-qPCR on the promoter regions of *ABCC3* and *ABCA4*. Compared with parental Daoy cells, V2-8 cells exhibited significantly greater enrichment of both p300 and the active histone mark H3K27ac at the *ABCC3* and *ABCA4* promoters (**Figure 6.A)** indicating enhanced transcriptional activation of these loci in resistant cells. Consistent with this, treatment with SGC-CBP30 markedly reduced H3K27ac enrichment at both promoters (**Figure 6.C**), supporting a role for the CBP/p300 bromodomain in maintaining their active chromatin state. At the transcriptional level, SGC-CBP30 treatment led to decrease in *ABCA4* expression in V2-8 cells, while such a prominent effect was not observed for *ABCC3* (**Figure 6.B**). Similar findings were observed in the independently generated V1-10 resistant model (**Figure S1.K**). On the other hand, A485 significantly reduced ABCC3 expression but increased *ABCA4* expression in both parental and resistant cells **(Figure S2.C-D)**, suggesting distinct contributions of the CBP/p300 bromodomain and HAT activities to the regulation of individual ABC transporters. Likewise, CRISPR-mediated depletion of CBP or p300 altered transporter expression, with CBP knockout significantly reducing *ABCC3* expression, whereas depletion of either of the genes decreased *ABCA4* levels (**Figure 6.D**).

**Figure 6.**
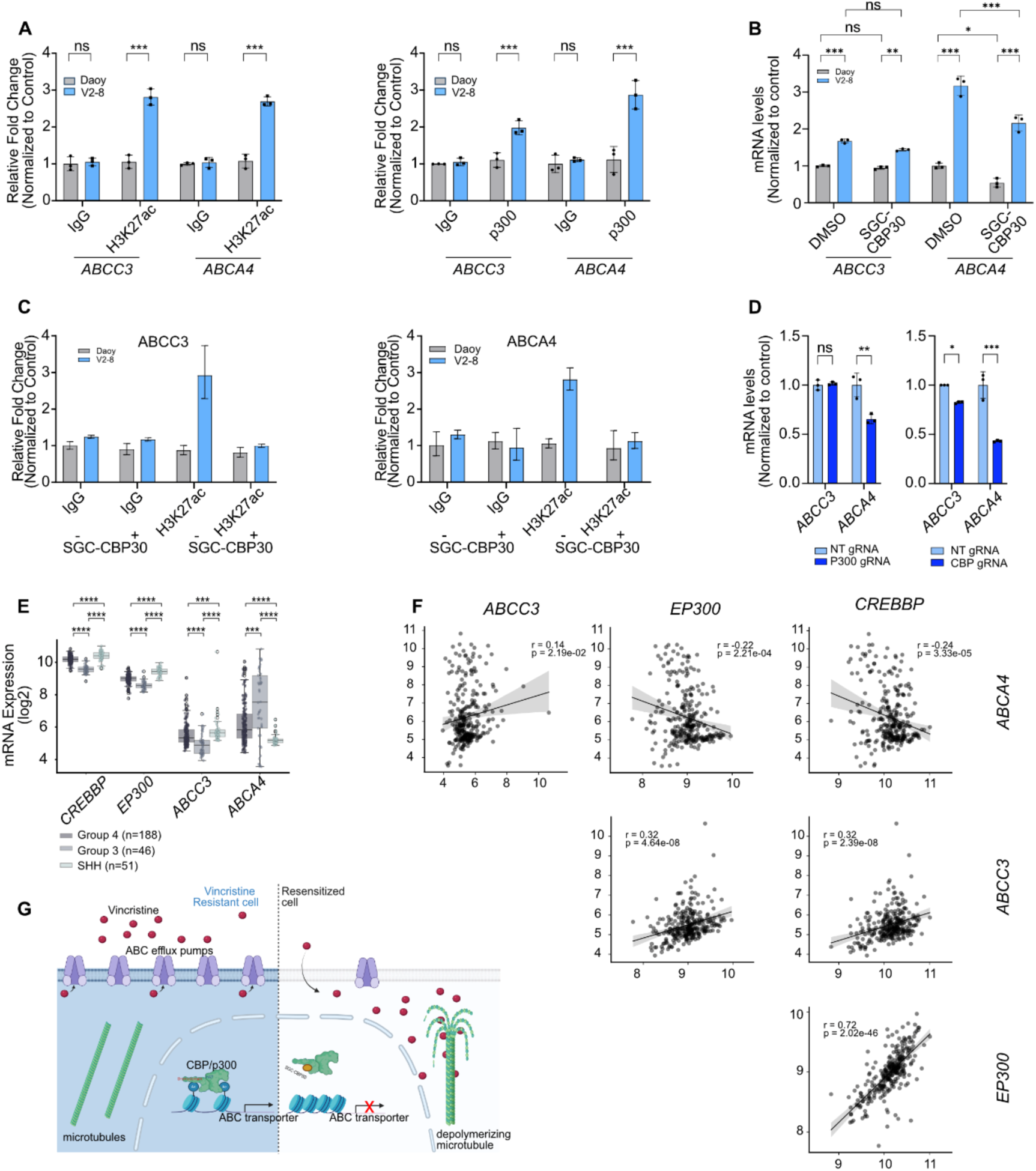
Acquired vincristine resistance is dependent on CBP/p300. A) CHIP-qPCR of H3K27ac histone mark and p300 on *ABCC3* and *ABCA4* in Daoy and V2-8 cells. B) *ABCC3* and *ABCA4* mRNA expression levels upon SGC-CBP30 treatment in V2-8 cells. C) CHIP-qPCR of H3K27ac marks on *ABCC3* and *ABCA4* in SGC-CBP30 treated Daoy and V2-8 cells. D) *ABCC3* and *ABCA4* mRNA expression levels upon knockout of respective genes in V2-8 cells. E) Expression of *CREBBP*, *EP300*, *ABCC3* and *ABCA4* on Northcott et. al. cohort in medulloblastoma subgroups. F) Correlation of *CREBBP*, *EP300*, *ABCC3* and *ABCA4* expression in Northcott et al cohort. G) Graphical model of results.

To evaluate the clinical relevance of this regulatory mechanism, we analyzed the Northcott MB cohort. *CREBBP*, *EP300*, and *ABCC3* expression were significantly elevated in the SHH subgroup compared with other molecular subgroups (**Figure 6.E**), consistent with the SHH classification of the DAOY cell line used in this study. Moreover, *ABCC3* expression positively correlated with both *CREBBP* and *EP300* across patient samples (**Figure 6.F**, **S.3**), supporting a clinically relevant CBP/p300–ABCC3 regulatory axis in MB **(Figure 6.G).**

## DISCUSSION

Vincristine is one of the most widely used chemotherapeutic agents in standard MB treatment regimens, yet its clinical benefit is frequently compromised by dose-limiting peripheral neuropathy and the development of acquired drug resistance^5^. Although recent clinical studies have explored vincristine-free treatment strategies for selected patients with previous vincristine-related toxicity^32^, vincristine continues to be incorporated into the majority of standard treatment regimens. Therefore, identifying therapeutic strategies that restore vincristine sensitivity rather than replace the drug remains an important clinical objective, particularly for relapsed disease.

In the present study, we identify CBP/p300 bromodomain activity as a previously unrecognized epigenetic regulator of acquired vincristine resistance in MB. Using independently generated resistant models, focused epigenetic compound screening and transcriptomic profiling, we demonstrate that pharmacological inhibition of the CBP/p300 bromodomain restores vincristine sensitivity through modulation of ABC transporter expression and activity. Together, these findings establish a clinically relevant mechanism linking chromatin regulation to chemotherapy resistance.

To identify targetable epigenetic dependencies associated with acquired vincristine resistance, we established two independently derived resistant Daoy cell lines using a dose-escalation strategy. Epigenetic compound screening revealed distinct chromatin dependencies between parental and resistant cells. Whereas broad-spectrum HDAC and HMT inhibitors induced cytotoxicity in both parental and resistant cells, consistent with previous reports^33,34^, bromodomain inhibitors emerged as the dominant class capable of selectively restoring vincristine sensitivity in resistant cells. Multiple bromodomain inhibitors targeting CBP/p300 and related bromodomain-containing proteins (CBP/BRD4, SGC-CBP30, I-CBP112, PFI-4, OF-1, and NVS-CECR2) displayed activity only in resistant cells when combined with vincristine, suggesting that acquired resistance creates a dependence on bromodomain-mediated transcriptional programs.

CBP and p300 are closely related lysine acetyltransferases that function as transcriptional coactivators through both their catalytic HAT domains and bromodomains, integrating chromatin acetylation with transcription factor recruitment and enhancer activity^24,35,36,37,38^. Although these proteins are best known for their roles in transcriptional regulation during development and tumorigenesis, increasing evidence indicates that their bromodomains independently regulate gene expression programs involved in cancer progression and therapeutic response^31,39,40,41^.

Previous studies have largely focused on BET family bromodomain inhibitors in MB, demonstrating that BRD4 inhibition suppresses subtype-specific oncogenic transcriptional programs, including GLI1 in SHH tumors and MYC in Group 3 disease^42,43^. Likewise, *CREBBP* has been identified as a recurrently altered gene in subsets of MB, supporting an important role for CBP/p300 biology in disease pathogenesis^44,45^. More recently, selective inhibition of the CBP/p300 bromodomain by SGC-CBP30 was shown to preferentially impair Group 3 MB through disruption of MYC-dependent transcriptional programs^46^. Our findings extend these observations by identifying a distinct function for CBP/p300 bromodomain activity in regulating acquired chemotherapy resistance. Notably, low-dose SGC-CBP30 substantially shifted the vincristine IC₅₀ of resistant cells, while higher concentrations restored sensitivity to levels approaching those of parental cells.

Our findings further support the utility of focused chemical probe libraries for identifying therapeutically actionable epigenetic vulnerabilities. Similar screening strategies have successfully uncovered novel dependencies across multiple cancer types including pediatric tumors, chordoma, pancreatic ductal adenocarcinoma, and breast cancer^47–50^. Importantly, the translational relevance of targeting the CBP/p300 bromodomain has recently been strengthened by the clinical development of inobrodib (CCS1477), a selective CBP/p300 bromodomain inhibitor derived from SGC-CBP30. Inobrodib has demonstrated clinical activity particularly in relapsed or refractory multiple myeloma, leading to fast-track designation by FDA^51,52^. Together with the data presented in this manuscript and ongoing development of new bromodomain inhibitors, these advances support CBP/p300 bromodomain targeting as a promising therapeutic strategy and provide a strong rationale for further studies on drug resistant medulloblastoma.

Mechanistically, our data support a model in which acquisition of vincristine resistance is accompanied by widespread transcriptional remodelling, including activation of multiple ABC and SLC transporter genes. Although *ABCB1*, the canonical vincristine transporter, represented one of the most highly upregulated genes in resistant cells, CBP/p300 inhibition preferentially affected *ABCC3* and *ABCA4*, suggesting selective regulation of a subset of transporter genes rather than global suppression of multidrug resistance pathways. ChIP-qPCR demonstrated increased p300 occupancy and H3K27 acetylation at the promoters of both genes in resistant cells, while bromodomain inhibition reduced promoter-associated H3K27ac, supporting direct chromatin regulation by CBP/p300.

Interestingly, however, ABCC3 and ABCA4 displayed distinct responses to different modes of CBP/p300 inhibition. Bromodomain inhibition with SGC-CBP30 reduced expression of both transporters, whereas inhibition of the catalytic HAT domain with A485 preferentially suppressed *ABCC3* while paradoxically increasing *ABCA4* expression. Similar differential regulation was observed in our validation experiments using the second resistant model. Despite these divergent transcriptional responses, both inhibitors restored vincristine sensitivity, suggesting that regulation of *ABCA4* is more complex than initially anticipated and may involve compensatory transcriptional mechanisms or additional chromatin-associated regulators beyond bromodomain-dependent chromatin recognition. In contrast, *ABCC3* consistently responded across pharmacological, chromatin, and functional assays, indicating that it likely represents the dominant downstream effector contributing to CBP/p300-mediated vincristine resistance.

Our findings also complement recent work demonstrating that the CBP/p300 bromodomain inhibitor I-CBP112 sensitizes triple-negative breast cancer cells to chemotherapy while suppressing ABC transporter expression^53^. Although both I-CBP112 and SGC-CBP30 target the CBP/p300 bromodomain, important pharmacological differences exist between these compounds, as I-CBP112 has been reported to paradoxically stimulate CBP/p300 acetyltransferase activity, whereas SGC-CBP30 lacks this property^54^. Nevertheless, the convergence of these independent studies suggests that bromodomain-dependent regulation of drug transporter programs may represent a broader mechanism of acquired chemotherapy resistance across multiple cancer types.

Importantly, analysis of patient transcriptomic datasets further supported the clinical relevance of our findings. Whereas *ABCC3* expression positively correlated with *CREBBP* and *EP300*, *ABCA4* displayed an inverse relationship. This observation closely mirrors our experimental findings, in which *ABCA4* responded differently to bromodomain and HAT inhibition, suggesting that although both genes exhibit CBP/p300 occupancy, only *ABCC3* consistently reflects the clinically relevant transcriptional program associated with CBP/p300 activity. Collectively, these findings identify the CBP/p300–ABCC3 axis as the more robust mechanism underlying acquired vincristine resistance in MB.

Several limitations should be acknowledged. Our study is primarily based on *in vitr*o models of acquired vincristine resistance, and future studies should determine whether this mechanism is conserved across additional MB subgroups, patient-derived models, and *in vivo* systems. Furthermore, although SGC-CBP30 is a well-characterized CBP/p300 bromodomain inhibitor, additional studies will be required to define the broader chromatin networks regulated by CBP/p300 during therapy adaptation. Finally, evaluation of pharmacokinetic properties, blood–brain barrier penetration, and toxicity profiles will be essential before clinical translation of CBP/p300 bromodomain inhibitors in pediatric brain tumors.

In conclusion, our study identifies CBP/p300 bromodomain activity as a key epigenetic regulator of acquired vincristine resistance in MB. By linking CBP/p300-dependent chromatin regulation to expression of drug transporter programs, particularly ABCC3, we provide a mechanistic rationale for targeting the CBP/p300 bromodomain to restore vincristine sensitivity and improve treatment responses in relapsed MB.

## Supporting information

AdditionalFile1

AdditionalFile2

AdditionalFile3

AdditionalFile4

AdditionalFile5

AdditionalFile6

AdditionalFile7

AdditionalFile8

AdditionalFile9

AdditionalFile10

AdditionalFile11

AdditionalFile12

## ETHICS APPROVAL AND CONSENT TO PARTICIPATE

Not applicable

## CONSENT FOR PUBLICATION

All authors have approved the manuscript for publication.

## AUTHORS’ CONTRIBUTIONS

G.K., O.Y.B., F.S.A., and T.L. contributed equally to this work. T.L. established the vincristine-resistant cell models and performed the initial epigenetic screen; F.S.A. performed screening validation, RNA-seq experiments, and prepared the initial manuscript preparation; G.K. conducted the bioinformatic analyses and mechanistic studies and wrote the manuscript; and O.Y.B. performed the major functional validation experiments and led the completion of the study. A.C.A., F.S.P., E.O.G. were involved in data generation. A.P.C. and U.O. contributed by providing the chemical library, supporting the sequencing experiments, and contributing to data interpretation. Approved final manuscript: all authors.

## ACKNOWLEDGEMENTS

We gratefully acknowledge the use of the services and facilities of the Koç University Research Center for Translational Medicine (KUTTAM), funded by the Presidency of Turkey, Head of Strategy and Budget. We thank James Dunford and Martin Philpott for their assistance with epigenetic probe library construction and RNA sequencing, and Ahmet Cingöz and Duygu Uçku for their technical help. Schematic figures were created with BioRender.com and licensed for publication (Fig 1.A, Fig. S1.A: LB29QVBC3H, Fig 1.H: UJ29QVBMX8, Fig 2: PS29QVBYZQ, Fig 3: ZA29QVCBF2, Fig 6.I: NZ29QVB2O5).

