## Supplementary material for "Targeting CBP/p300 Overcomes Acquired Vincristine Resistance in Medulloblastoma": Suplementary_info

### SUPPLEMENTARY FIGURES

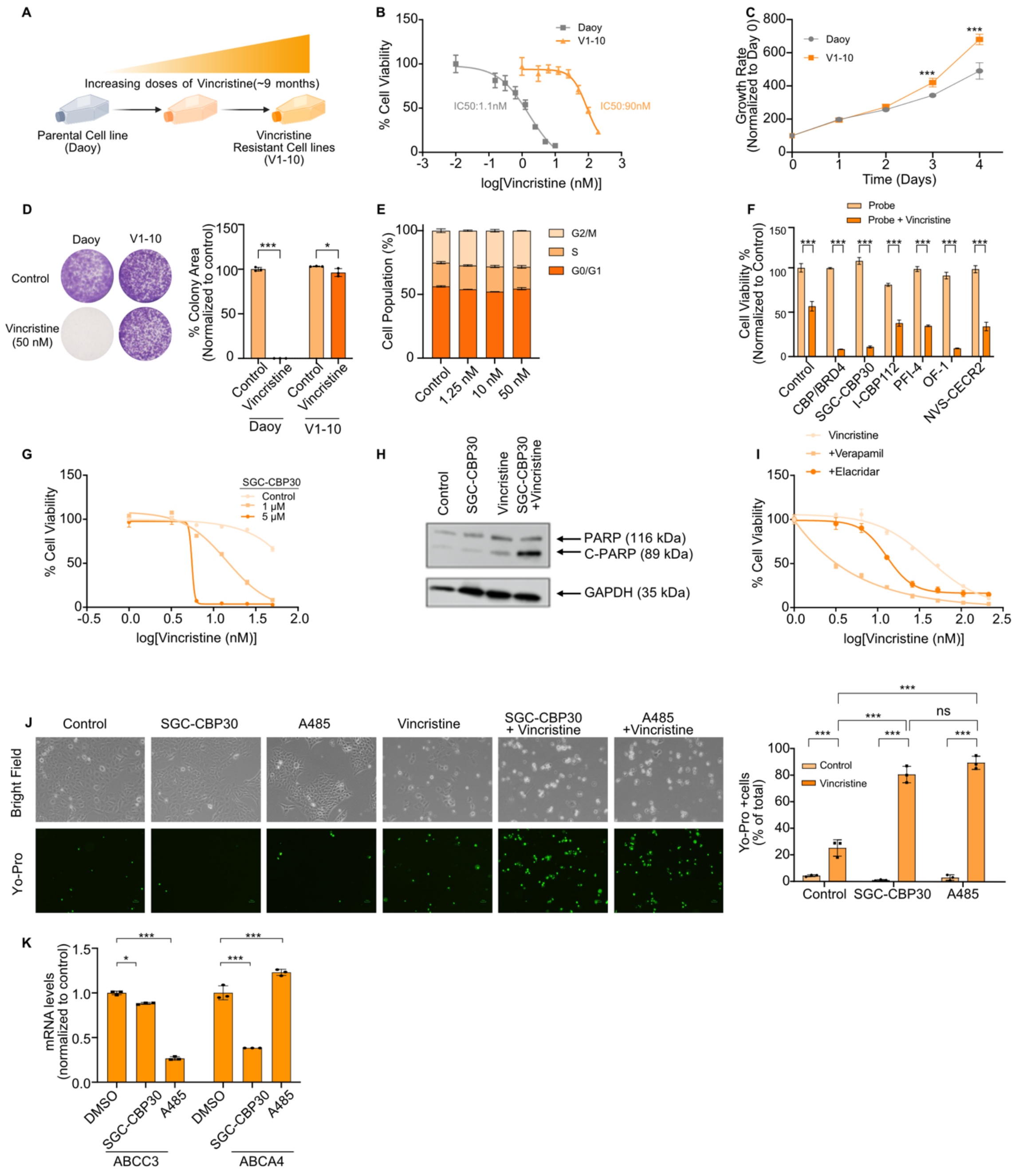


**Figure S1. Generation and characterization of V1-10 cell line.** A) Schematic representation of the establishment of V1-10 cell line using a dose-escalation approach. B) Dose-response curves of Daoy, and V1-10 cell lines treated with vincristine. C) Proliferation curves Daoy and V1-10 cell lines. D) Long-term clonogenic assay on Daoy and V1-10 cell lines treated with 50 nM vincristine. E) Cell cycle assay after 16 hours of DMSO or vincristine treatment (1.25, 10, 50 nM). F) Validation of selected bromodomain inhibitors in V1-10. G) Dose-response curves for vincristine and SGC-CBP30 combinations in V1-10. H) Western blot analysis of PARP cleavage in V1-10 cells treated with 1 nM or 50 nM Vincristine. I) Viability assay following treatment with Verapamil, Elacridar and/or vincristine for 72 hours. J) Yo-Pro staining of V1-10 cells treated with SGC-CBP30 in combination with vincristine to assess apoptotic cell death, K) ABCC3 and ABCA4 mRNA expression levels upon treatment with SGC-CBP30 or A485 on V1-10 cells.

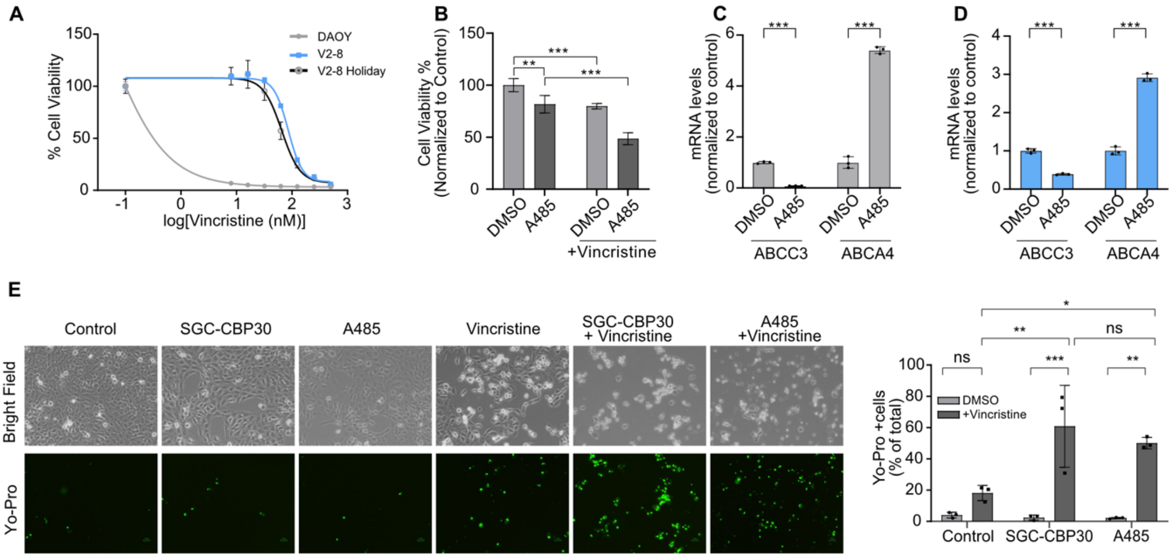


**Figure S2.** A) Dose-response curves of Daoy, V2-8 and V2-8-Holiday (cultured without vincristine for 50 days) cell lines treated with vincristine. B) Cell viability assay on Daoy andV2-8 cells treated with A485 and/or vincristine for 72 hours. *ABCC3* and *ABCA4* mRNA expression levels upon treatment with A485 on C) Daoy and D) V2-8 cells. E) YO-PRO-1 staining of Daoy cells treated with SGC-CBP30 in combination with vincristine to assess apoptotic cell death.

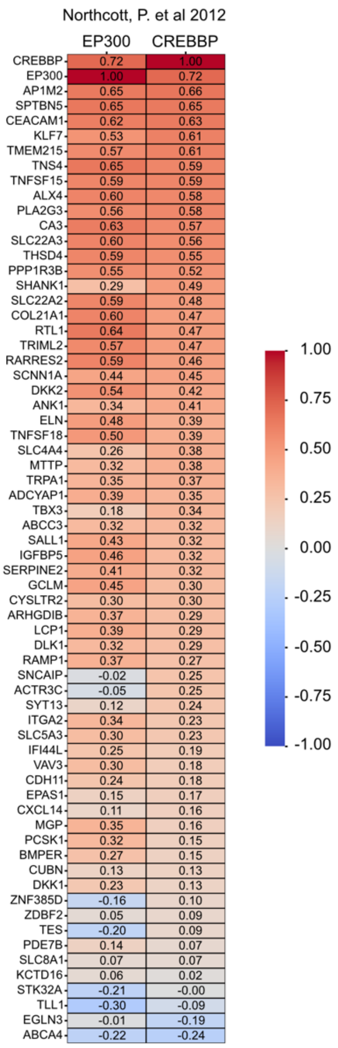


**Figure S3.** Correlation of the 69 genes with CREBBP and EP300 on cohort of Northcott et al.

### SUPPLEMENTARY TABLES

#### Supplementary Table 1: Composition of Chemical Probe Library and working concentrations of drugs in screens

| Probe | Class/Target | Working Concentration (µM) |
|---|---|---|
| (+)-JQ1 | Bromodomains - BRD2, BRD3, BRD4, BRDT (BET) | 1 |
| (-)-JQ1 | Bromodomains - Negative control | 1 |
| PFI-1 | Bromodomains - BRD2, BRD3, BRD4, BRDT (BET) | 5 |
| RVX-208 | Bromodomains - BRD2, BRD3, BRD4, BRDT (BET, BD2) | 5 |
| I-BET | Bromodomains - BRD2/3/4 | 1 |

| | | |
|---|---|---|
| Bromosporine | Bromodomains - pan-Bromodomain | 1 |
| CBP/BRD4 (0383) | Bromodomains - CBP, BRD4(1) | 5 |
| SGC-CBP30 | Bromodomains - CREBBP, EP300 | 1 |
| I-CBP112 | Bromodomains - CREBBP, EP300 | 1 |
| SMARCA | Bromodomains - SMARCA, PB1 | 2.5 |
| PFI-3 | Bromodomains - SMARCA2/4, PB1(5) | 1 |
| PFI-4 | Bromodomains - BRPF1B | 1 |
| OF-1 | Bromodomains - pan-BRPF | 5 |
| NI-57 | Bromodomains - pan-BRPF | 1 |
| BI-9564 | Bromodomains - BRD9, BRD7 | 1 |
| LP99 | Bromodomains - BRD9, BRD7 | 1 |
| I-BRD9 | Bromodomains - BRD9 | 10 |
| NVS-CECR2 | Bromodomains - CECR2 | 1 |
| BAZ2-ICR | Bromodomains - BAZ2A, BAZ2B | 1 |
| BAY-299 | Bromodomains - BRD1, TAF1 | 1 |
| Belinostat | HDAC - hydroxamic acids | 5 |
| SAHA | HDAC - hydroxamic acids | 2.5 |
| Trichostatin A | HDAC - hydroxamic acids - Class I & II | 0.5 |
| RGFP966 | HDAC - HDAC3 | 10 |
| PCI-34051 | HDAC - HDAC8 | 5 |
| Rocilinostat | HDAC - HDAC6 | 10 |
| Tubastatin A HCl | HDAC - HDAC6 | 10 |
| Valproic acid | HDAC - aliphatic acid compounds | 1000 |
| Entinostat | HDAC - ortho-amino anilides | 0.5 |
| SRT1720 | HDAC - SIRT1 (indirect?) activator | 1 |
| EX 527 | HDAC - SIRT1 | 1 |
| CI-994 | HDAC - 1,2,3,(8) | 1 |
| CXD101 | HDAC | 1 |
| PCI-24781 | HDAC - | 10 |
| Romidepsin | HDAC - | 1 |
| Mocetinostat | HDAC - | 10 |
| Santacruzamate | HDAC2? | 50 |
| CPI-360 | Histone methyltransferase - EZH2 and EZH1 | 10 |
| CPI-169 | Histone methyltransferase - EZH2, EZH1 | 10 |
| GSK343 | Histone methyltransferase - EZH2 | 3 |
| UNC1999 | Histone methyltransferase - EZH2 | 1 |
| UNC2400 | Histone methyltransferase - EZH2 | 1 |
| UNC0638 | Histone methyltransferase - G9a, GLP | 1 |
| UNC0642 | Histone methyltransferase - G9a, GLP | 1 |
| A-366 | Histone methyltransferase - G9a, GLP | 2 |
| LLY-507 | Histone methyltransferase - SMYD2 | 1 |
| BAY-598 | Histone methyltransferase - SMYD2 | 1 |

| | | |
|---|---|---|
| Chaetocin | Histone methyltransferase - SUV39H1 | 0.05 |
| A-196 | Histone methyltransferase - SUV420H1/H2 | 1 |
| PFI-2 | Histone methyltransferase - SETD7 | 2 |
| SGC0946 | Histone methyltransferase - DOT1L | 7.5 |
| Tranylcypromine | Lysine demethylases - LSD1 | 20 |
| GSK-LSD1 (irreversible) | Lysine demethylases - LSD1 | 0.5 |
| GSK-J4 | Lysine demethylases - JMJD3, UTX, JARID1B | 10 |
| GSK-J5 | Lysine demethylases - Negative control | 10 |
| IOX-1 | Lysine demethylases - pan-2-OG | 40 |
| KDOAM-25a | Lysine demethylases - JARID | 1 |
| KDOAM32 | Lysine demethylases - JARID | 1 |
| KDOPZ32 | Lysine demethylases - JARID | 1 |
| KDM5-C70 | Histone demethylase - JARID1 | 10 |
| Methylstat (Ester) | Histone demethylase | 2.5 |
| (E)-JIB-04 | Histone demethylase - Pan JmjC | 0.05 |
| ML324 | Histone demethylase - JMJD2E | 5 |
| KDOBA67 | Histone demethylase | 10 |
| OICR-9429 | Methyl Lysine Binder - WDR5 | 1 |
| UNC1215 | Methyl Lysine Binder - L3MBTL3 | 5 |
| A-395 | Methyl Lysine Binder - EED | 1 |
| A-395N | Methyl Lysine Binder - EED | 1 |
| SGC707 | Arginine methyltransferase - PRMT3 | 1 |
| TP-064 | Arginine methyltransferase - PRMT4 | 1 |
| TP-064N | Arginine methyltransferase - PRMT4 | 1 |
| MS049 | Arginine methyltransferase - PRMT4, PRMT6 | 1 |
| MS409N | Arginine methyltransferase - PRMT4, PRMT6 | 1 |
| GSK591 | Arginine methyltransferase - PRMT5 | 1 |
| MS023 | Arginine methyltransferase - Type I PRMTs | 1 |
| 5-Azacitidine | DNA methyltransferase (DNMT) - | 10 |
| 5-Azadeoxycitidine | DNA methyltransferase (DNMT) - DNMT1/3 | 5 |
| Olaparib | Poly ADP ribose polymerase (PARP) | 1 |
| Rucaparib | Poly ADP ribose polymerase (PARP) | 10 |
| 5-Iodotubercidin | Kinase inhibitor - ATP mimetic - Haspin | 1 |
| SGI-1776 | Kinase inhibitor - Haspin | 10 |
| CHR-6494 | Kinase inhibitor - Haspin | 1 |
| C646 | Histone acetyltransferase (HAT) p300/CBP | 1 |
| IOX2 | Prolyl-Hydroxylases - PHD2 (EGLN1) | 10 |
| GSK484 | Peptidyl arginine deiminase (PAD4) | 1 |
| GSK106 | Peptidyl arginine deiminase (PAD4) | 1 |
| GSK864 | Dehydrogenase | 5 |
| MAZ1805 | tRNA synthetase | 1 |

| | | |
|---|---|---|
| MAZ1392 | tRNA synthetase | 1 |

**Supplementary Table 2: Antibodies used in this study**

| Antibody | Purchased From | Dilution |
|---|---|---|
| PARP | Abcam, ab74290 | 1:2000 |
| GAPDH | Abcam, ab9485 | 1:2000 |
| α-tubulin | Cell Signaling (DM1A) | 1:10000 |
| Alexa Fluor® 488 Anti-IgG | Invitrogen | 1:500 |
| Goat anti-Mouse | Abcam, ab97023 | 1:10000 |
| Goat anti-Rabbit | Abcam, ab97051 | 1:10000 |

**Supplementary Table 3: qPCR primers used in this study**

| Transcript | Forward Primer (5'-3') | Reverse Primer (5'-3') |
|---|---|---|
| ABCA4 | TCGGGAACAGTCCCACGGAA | TTGAACACCCTTCACCGCCAA |
| ABCB1 | ACAGAGGGGATGGTCAGTGT | TCACGGCCATAGCGAATGTT |
| CREBBP | AGCAGCAGCTGGTTCTACTG | GGAGGCAAACAGGACAGTCA |
| GAPDH | AGCCACATCGCTCAGACAC | GCCCAATACGACCAAATCC |
| EP300 | GCAGTGTGCCAAACCAGATG | GGGTTTGCCGGGGGTACAATA |

**Supplementary Table 4: ChIP-qPCR primers used in this study**

| Transcript | Forward Primer (5'-3') | Reverse Primer (5'-3') |
|---|---|---|
| ChIP_ABCA4 | CATCTGAGACGCTGCACTAAC | TCTTCCAGCTACCAAACCCTTC |
| ChIP_ABCC3 | TCTACCTTCCCTCTTGGCGT | CCAGACTGCTACCACAAGGA |

**Supplementary Table 5: Sequences of sgRNAs used for CRISPR-KO experiment**

| Gene | Forward Sequence (5' to 3') | Reverse Sequence (5' to 3') |
|---|---|---|
| Non-targeting (NT) | GACGGAGGCTAAGCGTCGCAA | TTGCGACGCTTAGCCTCCGTC |
| CREBBP-gRNA | TGCCGGAAAGGTAATGACTC | GAGTCATTACCTTTCCGGCA |
| EP300 gRNA | CTGTAATAAGTGGCATCACG | CGTGATGCCACTTATTACAG |

**Additional Files**

| # | Filename | Fileformat | Title_of_data | Description |
|---|---|---|---|---|
| 1 | Additional File1 | m4v | Daoy_Control | Fig1D_Livecell |
| 2 | Additional File2 | m4v | Daoy_50nM | Fig1D_Livecell |
| 3 | Additional File3 | m4v | V2-8_Control | Fig1D_Livecell |
| 4 | Additional File4 | m4v | V2_8_50nM | Fig1D_Livecell |
| 5 | Additional File5 | m4v | Daoy_control | Fig3C_Livecell |
| 6 | Additional File6 | m4v | Daoy_CBP30 | Fig3C_Livecell |
| 7 | Additional File7 | m4v | Daoy_Vinc | Fig3C_Livecell |
| 8 | Additional File8 | m4v | Daoy_Combination | Fig3C_Livecell |

| | | | | |
|---|---|---|---|---|
| 9 | Additional File9 | m4v | V2-8_Control | Fig3C_Livecell |
| 10 | Additional File10 | m4v | V2-8_CBP30 | Fig3C_Livecell |
| 11 | Additional File11 | m4v | V2-8_Vinc | Fig3C_Livecell |
| 12 | Additional File12 | m4v | V2-8_Combination | Fig3C_Livecell |
